# Early Emergence of Perceptual Biases in Working Memory

**DOI:** 10.1101/2025.04.26.650749

**Authors:** Luis Serrano-Fernández, Ranulfo Romo, Néstor Parga

**Affiliations:** Departamento de Física Teórica, Universidad Autónoma de Madrid, 28049 Madrid, Spain; Centro de Investigación Avanzada en Física Fundamental, Universidad Autónoma de Madrid, 28049 Madrid, Spain; El Colegio Nacional, 06020 México DF, México

**Keywords:** Perceptual biases, Secondary Somatosensory cortex, Bayesian computations, Decision making, State-space geometry

## Abstract

Working memory (WM) is distributed across multiple cortical areas, suggesting that behaviors relying on WM arise from interactions between these regions. In a recent study, we found that during delayed comparison tasks, the first stimulus is not represented veridically in the prefrontal cortex (PFC), but instead is encoded in a systematically warped manner—biased toward the mean of the stimulus distribution. This neural distortion, which emerges already during the stimulus presentation and persists throughout the delay period, closely mirrors a contraction bias observed in behavior. Furthermore, the behavioral responses could be explained by a Bayesian observer model, in which the brain integrates prior expectations with noisy sensory inputs. These results suggest that the geometry of PFC neural trajectories embodies Bayesian estimates that underlie biased decisions. Here, we investigate whether the secondary somatosensory cortex (S2)—a lower-level sensory area also implicated in tactile WM—exhibits a similar encoding structure. Our analyses reveal that although WM-related signals in S2 are less robust than in PFC, the neural state space in S2 shares key geometric features with that of PFC, including a similarly warped representation of stimulus values. These findings suggest that perceptual biases may originate early in the cortical processing stream and are not exclusively shaped by higher-order associative areas. More broadly, our results support a distributed organization of WM, in which even sensory areas contribute to the formation of bias-prone representations that guide behavior.

## Introduction

Working memory (WM) enables the transient maintenance and manipulation of information no longer available in the sensory environment. Traditionally, higher-order associative regions such as the prefrontal cortex (PFC) have been considered the primary locus of WM operations [1], while sensory areas were thought to act mainly as transient relays. However, this view has evolved with increasing evidence that WM-related activity is distributed across multiple cortical areas [2]–[5]. Several studies have shown that sensory regions can maintain stimulus-related information during WM tasks, albeit with varying robustness [6].

One key insight from our recent work [7] is that WM-related population activity in PFC does not merely encode the stimulus veridically but reflects a systematic distortion observed in animal’s behavior: the value of the first stimulus in a delayed stimulus comparison task (a vibrotactile frequency, *f*_1_ [8]) is contracted toward the mean of the stimulus distribution. This perceptual bias, known as contraction bias or central tendency, has long been observed behaviorally in other tasks [9], [10]. In our previous study, we demonstrated that this bias can be quantitatively accounted for by a Bayesian observer model in which *f*_1_ is estimated as a compromise between the current sensory evidence and prior expectations based on stimulus statistics [7] (see also [11], [12]).

Importantly, we found that this contraction bias is evident in the geometry of PFC population activity during the presentation of *f*_1_, and persists throughout the subsequent delay period. Specifically, the relative distances between neural trajectories corresponding to different *f*_1_ values exhibit a sigmoidal dependence on stimulus value that mirrors the pattern of behavioral responses. The geometry of these neural trajectories closely matched the predictions of the Bayesian model, suggesting that PFC transforms sensory input into a behaviorally relevant estimate that incorporates prior knowledge. This transformation was not limited to WM maintenance but was also embedded in the initial encoding of the stimulus.

These findings raise fundamental questions about the broader cortical implementation of such biased representations. If PFC activity reflects a Bayesian estimate during the sensory encoding of *f*_1_, to what extent do other brain regions—particularly earlier sensory areas—contribute to this process? Is the bias already present in those regions, or does it emerge through transformations along the cortical hierarchy?

To address these questions, we focus here on the secondary somatosensory cortex (S2), a region involved in tactile perception that has also been implicated in WM maintenance [5], [6]. Although S2 lies lower in the cortical hierarchy than PFC, its neurons have been shown to exhibit selectivity to tactile frequencies and sustained activity during early stages of the delay period. We investigate whether S2 displays similar geometric signatures of Bayesian estimation and mnemonic distortion as those we previously observed in PFC.

Our analyses reveal that although WM signals in S2 are less robust than those in PFC, population activity in S2 shares key geometric features with PFC, including the warped, sigmoidal structure of stimulus representations. This suggests that the neural basis of perceptual biases may originate earlier in the cortical processing stream than previously assumed.

These findings support a distributed and hierarchical view of WM, where sensory areas contribute not only to the maintenance of stimulus information but also to the emergence of systematic Bayesian distortions in neural representations that are closely linked to behavioral biases.

## Results

### Tasks

This study analyzes neural recordings from S2 obtained in previous experiments [6] (Methods), in which two monkeys performed a vibrotactile frequency discrimination task (Fig. 1*A*). In this task, the animals had to compare two sequentially presented frequencies, *f*_1_ and *f*_2_, separated by a delay interval. After the presentation of *f*_2_, both monkeys were required to indicate their choice immediately. For one of the animals (monkey one), the duration of the delay period, Δ, varied across trials, taking one of five possible values: 1, 1.5, 2, 2.5 or 3 seconds. For the other animal (monkey two), the interval was fixed at 3 seconds for all trials. Moreover, the stimulus sets varied between the two monkeys. The first subject was exposed to set 1 (Fig. 1*B*, *Left*), while the second was presented with set 2 (Fig. 1*B*, *Right*). The probability distributions of *f*_1_ and *f*_2_ (Fig. 1*C*) are derived from these respective stimulus sets. Each set organizes the (*f*_1_, *f*_2_) frequency pairs (classes) into two diagonal groupings, referred to as the upper and lower diagonals. For clarity, each stimulus class was assigned a label, as indicated inside the squares in Fig. 1*B*.

**Fig. 1.**
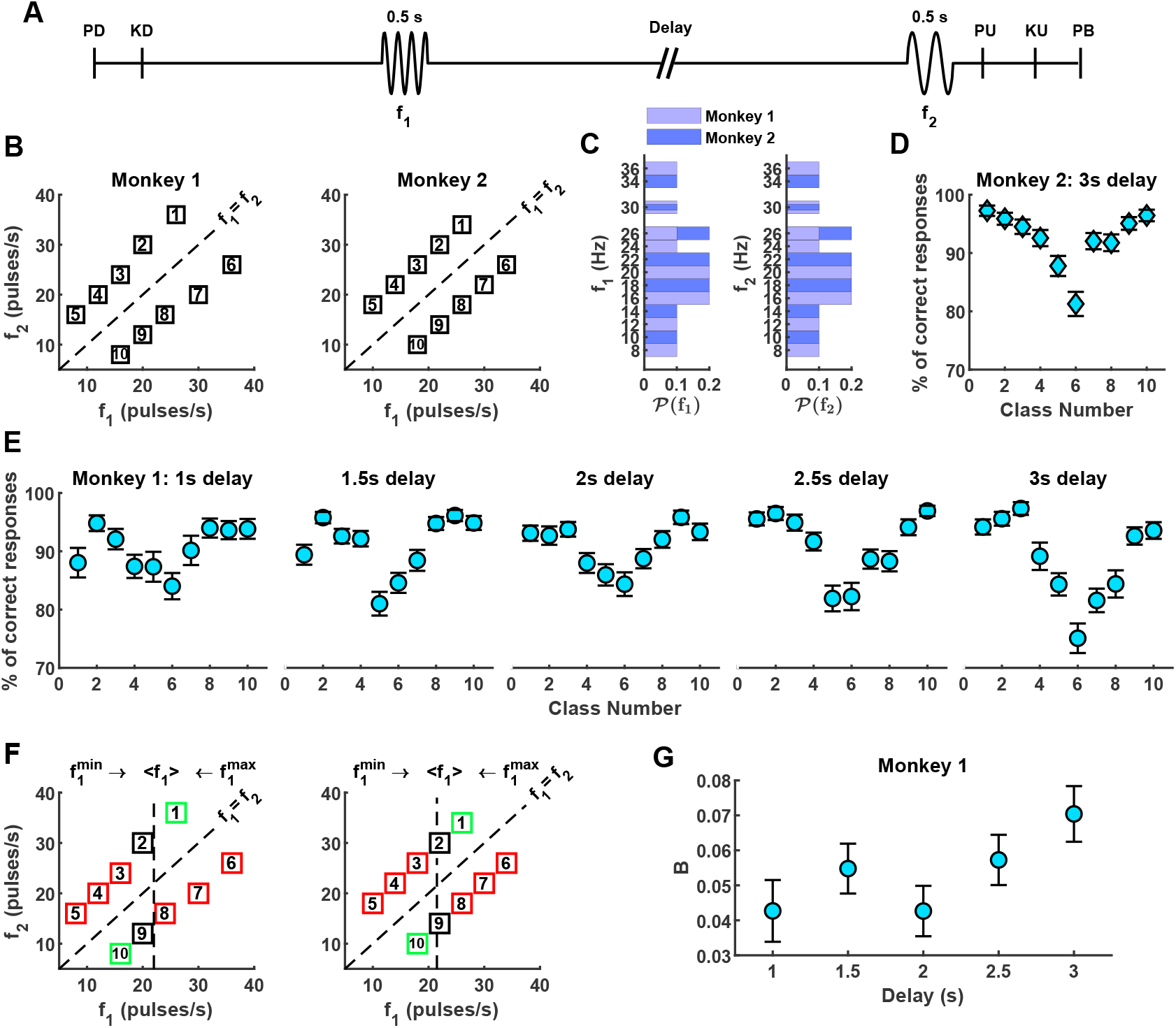
Task design, stimulus sets and behavior. *(A)* Timeline illustration of the frequency discrimination task performed by two monkeys. First, the smooth skin of a finger on the monkey’s restrained hand is contacted by a mechanical probe (Probe Down, PD). To begin the trial, a key is pressed (Key Down, KD) by the monkey with its unrestrained hand. Then, a 500 ms mechanical vibration (first stimulus, *f*_1_) is then delivered, followed by a delay period. For monkey one, the delay duration varied from trial to trial, with possible values of 1, 1.5, 2, 2.5 or 3 seconds. For monkey two, the delay was fixed at 3 seconds. Subsequent to the delay period, the second stimulus (*f*_2_) is applied as a 500 ms mechanical vibration. Once the probe is lifted (Probe Up, PU), the monkey uses its free hand (Key Up, KU) to press one of two push-buttons (PB), reporting its choice in the comparison between the two stimuli. *(B)* Sets of possible frequency pairs (classes: *f*_1_, *f*_2_) for monkey one (set 1; *(Left)*) and monkey two (set 2; *(Right)*). *(C) 𝒫* (*f*_1_) and 𝒫 (*f*_2_) indicate the probability of the first *(Left)* and second *(Right)* frequency, respectively, according to the stimulus sets used for monkeys one (light purple) and two (dark blue). *(D)* Percentage of correct trials for monkey two as a function of stimulus class. *(E)* Percentage of correct responses for monkey one. From left to right, each panel corresponds to one of the five possible delay durations (1s, 1.5s, 2s, 2.5s, and 3s). *(F)* Due to contraction bias, the first stimulus (*f*_1_) is perceived as contracted towards its distribution mean, increasing task difficulty for red classes and facilitating performance for green classes. *Left* panel corresponds to set 1 (monkey one), while *right* panel corresponds to set 2 (monkey two). *(G)* Contraction bias is quantified as the root-mean-square deviation of performance (*B*; Eq. 8). Error bars computed with 1,000 bootstrap resamples.

For the remainder of this study, we begin by analyzing the behavioral data, followed by evaluating its fit to a behavioral model. We then examine neural activity in the S2, focusing on its geometry in state space and how and to what extent this region maintains the value of the first stimulus through its firing patterns. Finally, we explore the relationship between behavior and population-level neural dynamics.

### Behavior. Contraction bias shaped monkeys’ performance

Since the difference between the two frequencies is constant across classes in set 2 (Δ*f* = |*f*_1_ − *f*_2_| = 8Hz), or takes one of two absolute values in set 1 (Δ*f* = 8Hz or 10Hz), the task difficulty should be comparable across classes, and performance is expected to be similar. However, the percentage of correct responses exhibited class-dependent variations (Fig. 1*D* for monkey two and Fig. 1*E* for monkey one, with one panel per delay duration). These class-dependent variations can be attributed to the contraction bias [7], [9], [11]–[13], which differentially affected decision accuracy depending on the specific pairing of stimuli. For the two monkeys, this effect induced a shift in the perceived frequency of the first stimulus, *f*_1_, toward the mean of its frequency distribution (i.e., ⟨*f*_1_⟩ = 24Hz and ⟨*f*_1_⟩ = 22Hz, as derived from the light purple and dark blue distributions, respectively, in Fig. 1*C*, *Left*). Consequently, lower values of *f*_1_ tended to be perceived as higher, while higher values were perceived as lower (see titles of Fig. 1*F*). Thus, the influence of bias progressively impaired the accuracy for classes where *f*_1_ *< f*_2_ (upper diagonals in Fig. 1*B*), encountered from right to left. Conversely, it increasingly facilitated performance for classes where *f*_1_ *> f*_2_ (lower diagonals). To preserve the bias structure, we labeled classes at the extremes of the sets as those benefiting from the bias (green in Fig. 1*F*), while the central classes were those negatively affected by its effects (red in Fig. 1*F*). As previously observed in [7], [11], [12], plotting accuracy against this class labeling revealed V-shaped patterns (see Figs. 1*D* and *E*), as a consequence of the contraction bias. These behavioral patterns raise the question of what underlying computational mechanisms might give rise to the contraction bias. The next section outlines our rationale for focusing on how prior knowledge, combined with current-trial observations, may give rise to this bias, in line with Bayesian principles.

### Bayesian Influences on Monkey Behavior

Contraction bias was quantified as the root-mean-square (RMS) deviation of the animal’s performance, *B* (see Methods). For monkey two, who performed the task with a fixed 3-second delay, the quantified bias was *B* = 0.0443 ± 0.0054. On the other hand, since monkey one performed the task with variable delay durations, *B* was calculated separately for each delay condition (Fig. 1*G*). Here, *B* increases with the duration of the delay interval. This increase in bias magnitude is consistent with predictions from the Bayesian framework, which suggests that maintaining *f*_1_ in short-term memory leads to greater uncertainty about its value [7], [12], [14]. This is why we focus on how prior knowledge, combined with current-trial observations, can give rise to contraction bias through the formulation of a normative model.

### Bayesian Model

To analyze the behavioral data of the two monkeys, we applied the same Bayesian model used in our recent study on PFC neuron recordings during the tactile discrimination task [7]. Below, before applying the model to the current data, we summarize the key features of the model. We considered an observer who makes imperfect measurements of the stimulus frequencies *f*_1_ and *f*_2_, has prior knowledge of the stimulus set—including its mean and variance—and forms beliefs based on these observations. In our model, we assumed that frequency measurements follow a Gaussian distribution, with means *f*_1_ and *f*_2_, and standard deviations defined as *σ*_1_ = *W*_1_*f*_1_ and *σ*_2_ = *W*_2_*f*_2_, where *W*_1_ and *W*_2_ correspond to the respective Weber fractions. Furthermore, we assumed that the long-term sensory history of the first frequency plays a role in shaping perceptual decisions. Prior models of contraction bias have incorporated dependencies on the mean of the stimulus distribution [12], [13], [15], [16], and experimental findings suggest that such dependencies align with a Bayesian framework [7], [17].

To model decision-making in the current trial *n*, we proposed that the observer’s judgment is influenced by a combination of the noisy measurement of the first frequency 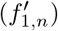 and an estimate of its stationary sensory history. Formally, we expressed the observation *o*_1,*n*_ as:

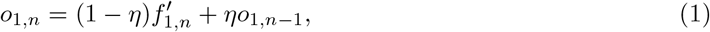

where *η* = 0 implies that long-term sensory history has no influence, whereas *η* = 1 indicates that the decision relies entirely on past observations, disregarding the current stimulus. In the Gaussian approximation, the likelihood *P* (O_1_|*f*_1,*n*_) represents the probability of the stochastic Gaussian variable O_1_—which corresponds to the observed stimulus in the current trial—given that the presented first frequency is *f*_1,*n*_. The mean and variance of this likelihood function are given by [7]

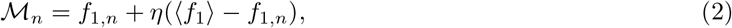

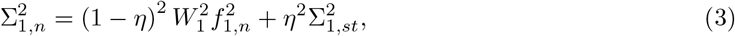

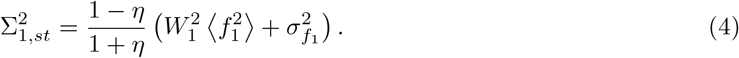

Here, 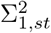 represents the stationary variance of the first frequency’s observations. The terms 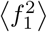 and 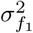 denote the second moment and variance of the first frequency within the stimulus distribution.

Importantly, both ℳ and Σ_1_ contribute to the contraction bias effect. The mean ℳ_*n*_ explicitly reflects the tendency of *f*_1,*n*_ to shift toward ⟨*f*_1_⟩, with the magnitude of this shift being proportional to the difference between the actual frequency and the distribution mean. Bayesian influence on contraction bias arises when the prior exerts a stronger effect on the posterior of the first frequency than on the second, particularly in cases where Σ_1_ is greater than *σ*_2_ [7], [9], [12]. Priors correspond to the probability distribution of stimuli (Fig. 1*C*) and encode the underlying structure of the stimulus set (Methods). Lastly, we assume that *f*_2_ is affected only by measurement noise, with variance 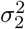.

At the end of the presentation of the first frequency, the observer forms a belief about its value, denoted as 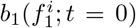, with components on all possible values of *f*_1_ within the stimulus set (*i* = 1, …, *N*_1_) (Fig. 1*B*). This belief is based on Bayes’ theorem

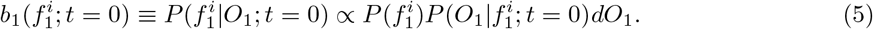

Note that the onset of the delay period (the end of the presentation epoch) was taken as the time origin. The left-hand side indicates that the posterior probability defines 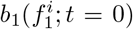.Because the comparison between the two frequencies occurs during the presentation of *f*_2_, the belief about the first frequency must be propagated until the end of the memory period. To achieve this, we updated the belief about *f*_1_ at various times throughout the delay period. The temporal evolution of this belief can be recursively computed under the assumption of the Markov property [18]

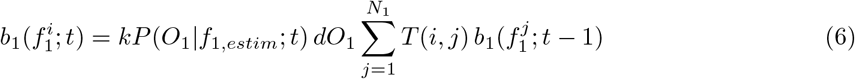

where the initial value of the belief is given by Eq. 5, *N*_1_ represents the number of possible values for the first frequency in the set and *k* is a normalization constant. *T* (*i, j*) is the probability of transitioning from state 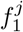 at time *t* − 1 to 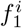 at time *t*. Given the assumption that the agent has good knowledge of the stimulus set, we took *T* (*i, j*) = *δ*_*i,j*_, where *δ*_*i,j*_ is the Kronecker delta, ensuring that the state remains unchanged across time steps. In order to propagate the belief throughout the delay period, during which the representation of the first stimulus is maintained in WM somewhere in the brain, we assumed that several estimates *f*_1,estim_(*t*) are made during the delay period (details about the estimators can be found in the Methods section).

At the end of the delay period (*t* = Δ), when the second frequency is presented, its posterior probability 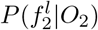, or belief 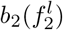, combines its prior with the noisy observation *O*_2_

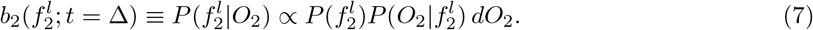

Here 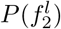 is the discrete prior probability of the second frequency (Fig. 1*C*, *Right*) and *l* labels the possible values of that frequency in the stimulus set (Fig. 1*B*). Note that we should specify whether this prior probability depends on the presented value of the first frequency. We assumed that the noise affecting that frequency is sufficiently large, making this dependence minimal by the time the second stimulus is presented.

At this stage, the two beliefs are compared to determine which of the two frequencies was larger. It is important to note that the observations of the two frequencies were made during their respective presentation epochs. Specifically, 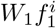 represents the standard deviation of the measurement noise for 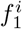 before the delay period, while 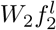 corresponds to the standard deviation of the measurement noise for 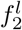 at the end of the delay period. Consequently, by the time the comparison between stimuli takes place, *W*_1_ and *W*_2_ are expected to differ. In [7], where the delay duration was fixed, both parameters were defined at the end of the delay period, since belief propagation throughout the delay had not been included in the model.

### Fits of Behavioral Data

To estimate the model parameters (*η, W*_1_, *W*_2_; see Table 1), we analyzed the accuracy curve specific to each monkey (Fig. 1*D,E*) and leveraged it to quantify the similarity between the model’s predictions and the experimental data (Methods). To fit the model, in Eq.6 we assumed that the animal estimated *f*_1_ during the delay period using a Bayesian estimator, that is, we took *f*_1,*estim*_ as *f*_1,*Bayes*_ (see Methods, Eqs. 11-12; we will return to this assumption later). Since data for monkey one were collected across multiple durations of the memory epoch, we fitted the accuracy curve separately for each of these durations (Fig. 2*A*). The corresponding bias values (*B*) were calculated as a function of Δ from both the observed and fitted accuracy curves (Fig. 2*B*). An analogous fitting procedure was applied to the accuracy curve of monkey two (Fig. 2*C*), for which *B* was 0.0443 ± 0.005 vs 0.044. For both monkeys, the belief about the first stimulus was propagated throughout the delay period, dividing each delay duration Δ into 500 ms subintervals.

**Table 1.**
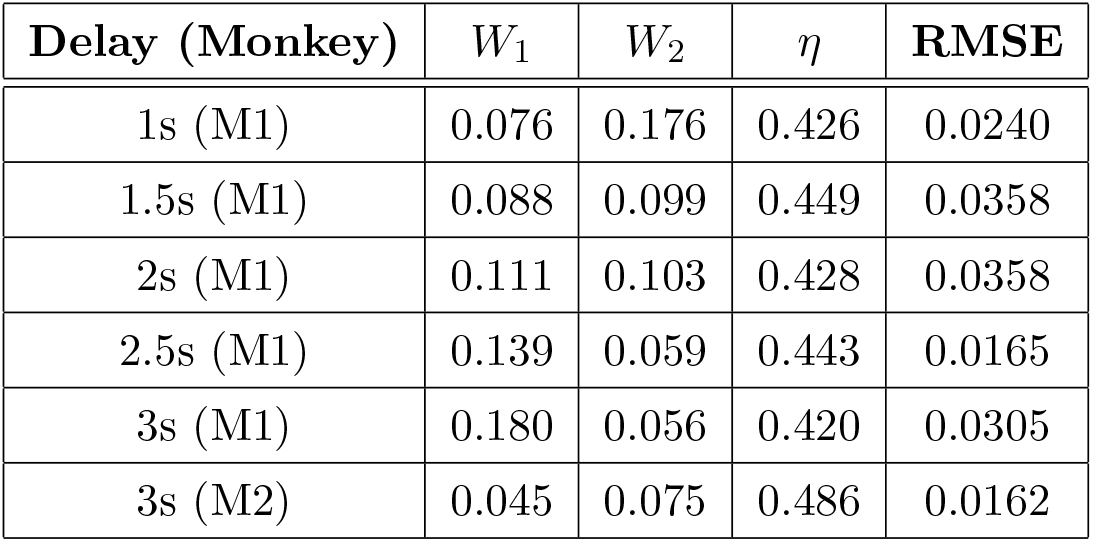
Bayesian model fits across delay conditions for two monkeys. The fitted behavioral datasets correspond to the five delay conditions tested in monkey one (1, 1.5, 2, 2.5 and 3 seconds), and the 3-second delay condition in monkey two. The second and third columns display the fitted measurement uncertainty parameters (*W*_1_ and *W*_2_). The fourth column shows the fitted sensory history parameter (*η*), and the last column reports the root mean square error (RMSE) between behavioral and model performance.

| **Delay (Monkey)** | $W_1$ | $W_2$ | $\eta$ | **RMSE** |
| --- | --- | --- | --- | --- |
| 1s (M1) | 0.076 | 0.176 | 0.426 | 0.0240 |
| 1.5s (M1) | 0.088 | 0.099 | 0.449 | 0.0358 |
| 2s (M1) | 0.111 | 0.103 | 0.428 | 0.0358 |
| 2.5s (M1) | 0.139 | 0.059 | 0.443 | 0.0165 |
| 3s (M1) | 0.180 | 0.056 | 0.420 | 0.0305 |
| 3s (M2) | 0.045 | 0.075 | 0.486 | 0.0162 |

**Fig. 2.**
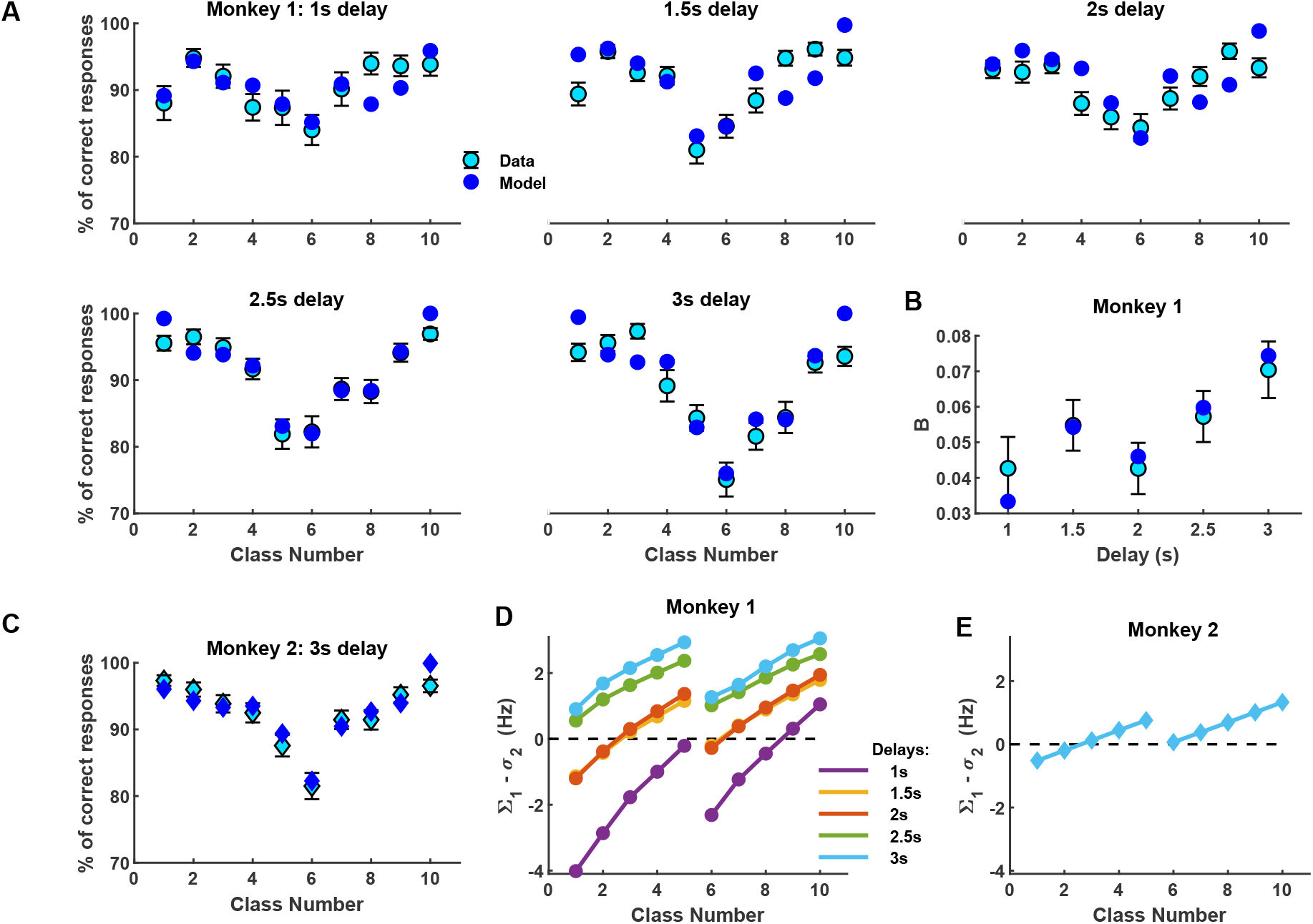
Bayesian fits of the accuracy curves. *(A)* Fits for monkey one. Each panel corresponds to one of the five possible delay durations (1s, 1.5s, 2s, 2.5s and 3s). Cyan circles with error bars correspond to behavioral data, and dark blue circles represent the Bayesian fit. Error bars computed with 1,000 bootstrap resamples. *(B)* Measurement of contraction bias (*B*; Eq. 8) obtained from behavioral performance (cyan markers) and the Bayesian fit (dark blue markers). *(C)* Bayesian fit for monkey two. Cyan diamonds with error bars correspond to behavioral data, and dark blue diamonds represent the Bayesian fit. *(D)* Bayesian contribution to the bias vs. class number for several values of the delay period for monkey one. This contribution is determined by the quantity Σ_1_ − *σ*_2_, where Σ_1_ is given by Eq. 8 and *σ*_2_ = *W*_2_*f*_2_. As delay duration increases, Σ_1_ − *σ*_2_ becomes increasingly positive, reinforcing the Bayesian contribution to the bias. *(E)* Bayesian contribution to the bias for monkey two. The majority of classes (*f*_1_, *f*_2_) satisfy the condition Σ_1_ *> σ*_2_.

Next, we examined the contributions of stimulus uncertainty and stationary sensory history to the contraction bias. Bayesian contribution of the bias emerges when the prior has a stronger influence on the posterior of the first stimulus than on that of the second [9]. This is especially evident when the uncertainty associated with the first frequency (Σ_1_(*f*_1_; *t* = Δ)) is greater than that of the second (*σ*_2_). For monkey one (Fig. 2*D*), the difference Σ_1_(*f*_1_; *t* = Δ) − *σ*_2_, evaluated at the end of the delay period, closely paralleled the empirical relationship between *B* and Δ (Fig. 2*B*). Specifically, for the shortest delay duration, the uncertainty difference was negative for most stimulus classes; however, as Δ increased, this pattern changed, with the difference becoming positive for the majority of stimulus classes (Fig. 2*D*). This finding again suggests that longer delay periods enhance the influence of Bayesian inference on the contraction bias. In contrast, as expected the sensory history parameter *η* remained largely unchanged as the delay interval increased (Table 1). For monkey two, we could not directly test this Bayesian prediction because only one delay interval was used in the experiment.

Nevertheless, analysis of the stimulus uncertainties (Fig. 2*E*) revealed that Bayesian inference likely contributes significantly to the bias, as the uncertainty difference was positive for all stimulus classes except classes one and two.

### S2 Neurons Carry Information About the Current but Not the Previous Trial’s *f*_1_

Sensory history has been suggested as a source of contraction bias [12], [13], [19], potentially shaping perception through persistent neural activity from past stimuli or by integrating inputs from other brain regions. However, since contraction bias likely emerges as a network-level phenomenon spanning multiple areas [7], [13], [17], the presence of such bias does not necessarily imply that individual neurons in a given region encode past sensory experiences. For instance, neurons in the monkey PFC, a key region for WM, contribute to contraction bias through Bayesian computations [7] yet do not encode short-term sensory history. Given the distributed nature of WM [2], [5], information about prior stimuli may instead be stored elsewhere.

The secondary somatosensory cortex is the first cortical region where mnemonic neural activity related to the current trial *f*_1_ is observed, though only during the early delay period [5]. This activity re-emerges at the onset of the second stimulus, and S2 plays a role in comparing *f*_1_ and *f*_2_ [6]. If these neurons were influenced by recent stimulus history, this effect could contribute to the behavioral bias observed in the animals. To test this, we quantified the MI each neuron carried about both the current trial *f*_1_ and its value from the preceding trial (Methods). In agreement with previous analyses [20], a subset of S2 neurons in monkey one exhibited significant MI about the current value of *f*_1_ during its presentation and extending into the early phase of the delay period (Fig. 3, blue line; see also SI Appendix, Fig. S2, and three example neurons in Fig. S3). However, this MI signal disappeared as the delay progressed. Interestingly, at the onset of the second stimulus, MI about *f*_1_ re-emerged and remained detectable even after *f*_2_ presentation [20]. A central question is whether S2 neurons also carried MI about the first stimulus from the preceding trial. Our analysis revealed no evidence of such information (Fig. 3, red line). An analogous analysis was conducted for monkey two. While similar patterns of MI were observed—both with respect to the current and previous *f*_1_—the effects were weaker in this animal (see SI Appendix, Figs. S4 and S5). A similar result had been found in PFC neurons recorded in the same task [7].

**Fig. 3.**
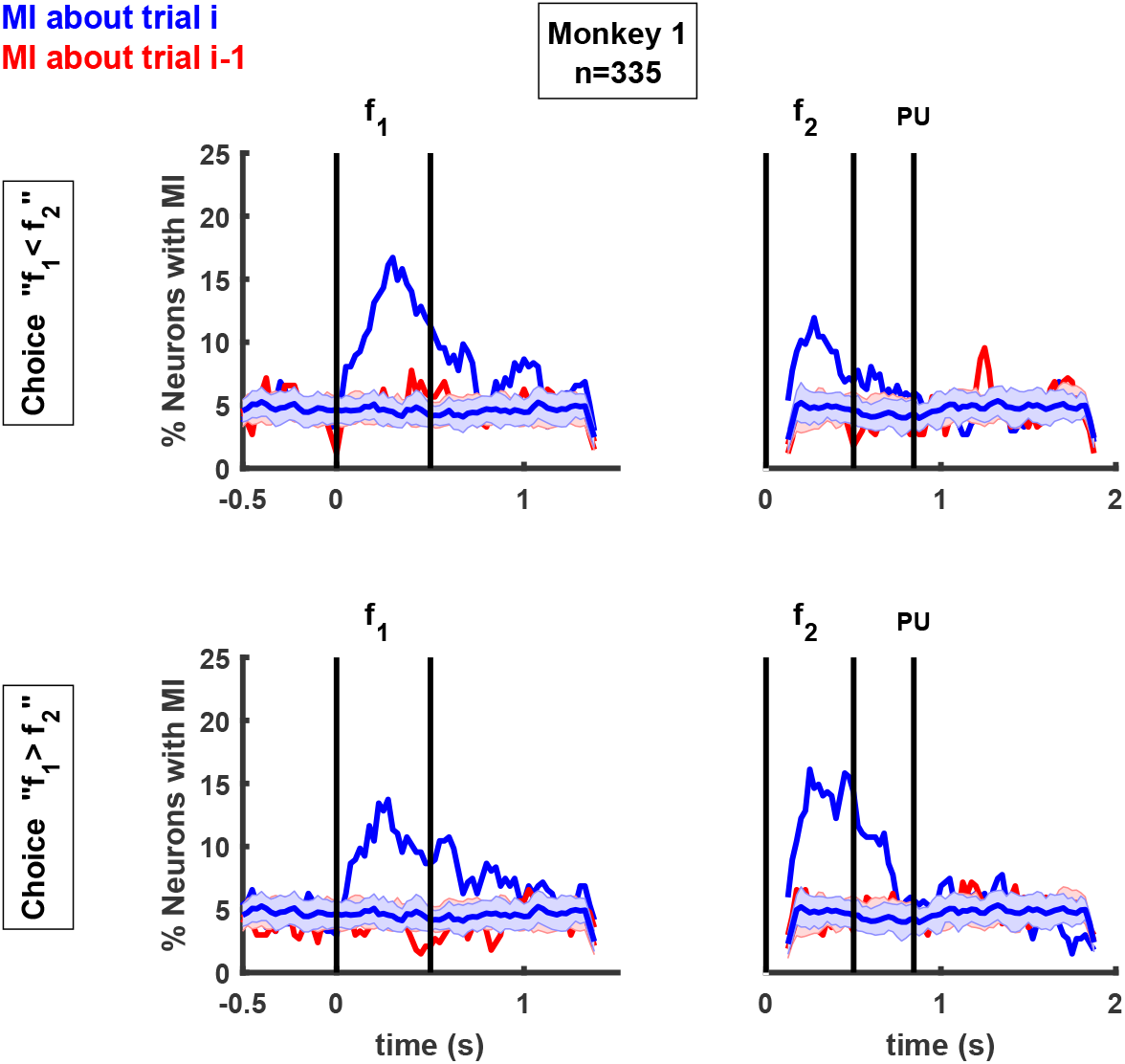
S2 neurons from Monkey one exhibit significant mutual information (MI) about current-trial *f*_1_, but not about the previous-trial *f*_1_. Percentage of neurons showing significant MI with respect to the first stimulus of the current trial (blue line) versus that of the previous trial (red line). The analysis was restricted to correct trials in which the current choice was fixed (*Top*: “*f*_1_ *< f*_2_”; *Bottom*: “*f*_1_ *> f*_2_”). Note that this constraint effectively fixes the stimulus class in the current trial. Neural recordings from all delay conditions were pooled to analyze MI, aligning activity to the onset of the first stimulus (left panels) or the second stimulus (right panels). Horizontal blue and red shadings indicate the mean and standard deviation computed from shuffled data for the current and previous trial, respectively (see Methods).

These findings specifically pertain to transient sensory history and do not exclude the possibility that PFC neurons encode information on the long-term stationary stimulation history.

### State-space analysis

Previous analyses of WM in this dataset have focused primarily on the activity of single neurons. Similarly, the MI evaluations presented here and in [20] have been based on single-cell responses. However, neural activity at the population level may exhibit strong structure and organization, revealing properties that may not be apparent from single-neuron analyses alone. Population activity is confined to a low-dimensional subspace within the high-dimensional neural state space [7], [12], [21], [22]. Within this subspace, neural activity trajectories during the delay period form structured manifolds whose geometry encodes stimulus parameters, providing a population-level representation that is more robust than that of individual neurons [7]. These findings suggest that to fully understand WM and decision-making mechanisms, it is essential to consider the collective geometry and dynamics of population activity. We then focused our analysis on neural trajectories during both the presentation of the first stimulus and the subsequent delay period. In correct trials, population neural activity was projected onto the first three principal components (Fig. 4), giving rise to neural trajectories corresponding to each *f*_1_ value that tend to evolve separately throughout the presentation of the first stimulus (Fig. 4*A,E*). This separation between trajectories can be observed in the relative distances with respect to a reference trajectory (Fig. 4*B,F*; see Methods). Additionally, state-space trajectories during the early phase of the delay (Fig. 4*C,G*) maintained that separation across *f*_1_ values (Fig. 4*D,H*). However, as the delay period progressed, the trajectories gradually lost their initial organization. To assess the extent and dynamics of WM deterioration over time, we extended the analysis to a higher-dimensional state space that captured a larger proportion of the variance (Fig. 5). In particular, during the presentation of *f*_1_, we constructed a subspace that explained at least 85% of the total variance (first column in Fig. 5), which gave rise to average relative distances with a sigmoidal dependence on the value of the first frequency. Continuing with the delay period, we investigated how the sigmoidal structure of the averaged relative distances evolved over time. To this end, we segmented the interval into half-second epochs and evaluated the temporal mean of the relative distances within each segment. In the first half-second, around 61% of the total variance was captured within 6–7 dimensions, and the sigmoidal pattern in the distances persisted (second column in Fig. 5). In the subsequent half-second, explaining the same proportion of variance required 11 dimensions, indicating a substantial expansion of the manifold defined by the trajectories (third column). Despite this increase in dimensionality, the relative distances maintained a similar pattern to the previous interval. In the following second, the dimensionality increased only slightly to 12 and the relative distances became noisier, suggesting that the trajectories exhibited less structured relationships (fourth and fifth columns). Note that examining the sequence of panels within the same row allows for a clear visualization of this effect within the same data. Thus, contrary to what the MI analysis suggested, the information encoded within the neural manifold did not collapse entirely during the delay. Instead, while representations became less distinct, some aspects of the stimulus-related structure remained visible in the population activity.

**Fig. 4.**
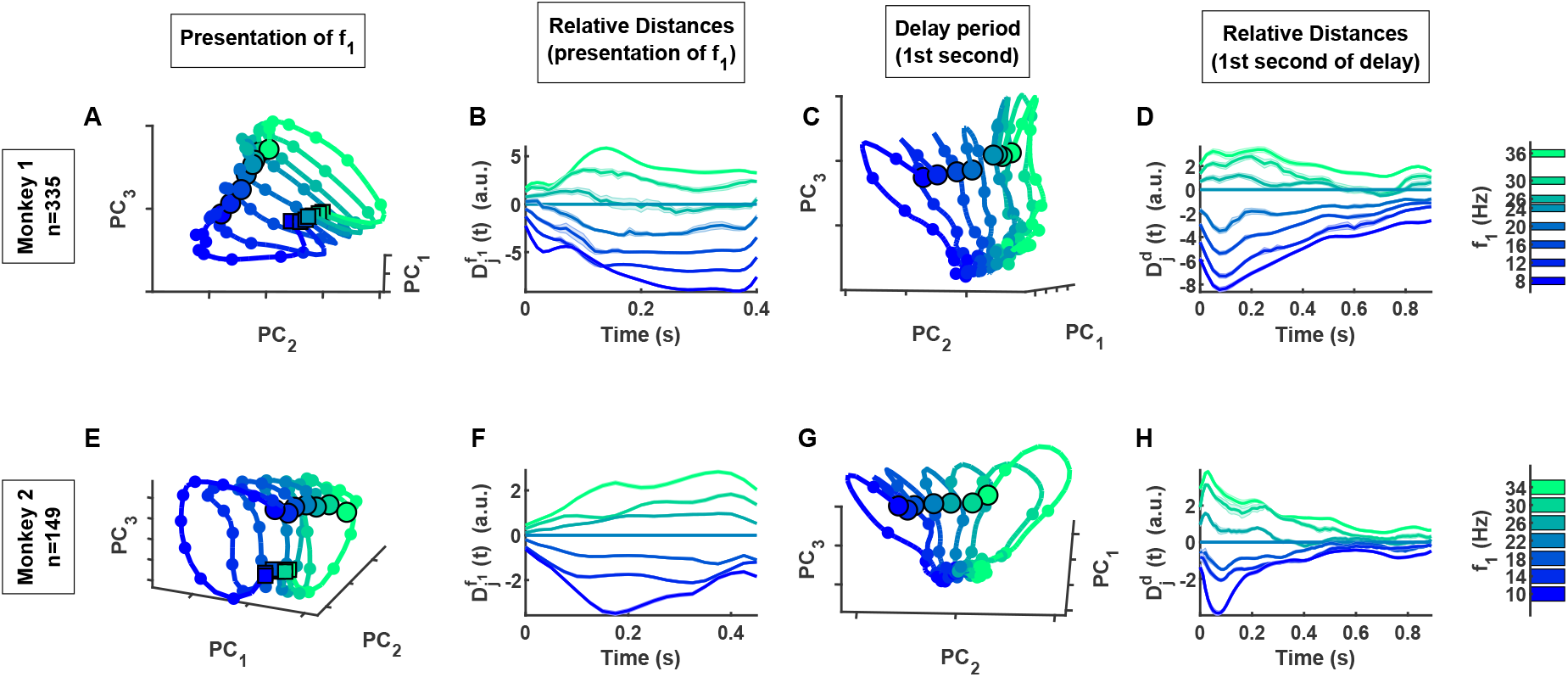
State-space trajectories and their relative distances. *(A–D)* Monkey one. Data from all delay conditions were pooled to analyze state-space activity during the presentation of *f*_1_ and throughout the delay period up to the end of its first second. *(A)* Population neural activity projected onto the first three PCs (61% of variance) during the presentation of the first stimulus. The onset and offset of *f*_1_ are indicated by squares and large circles, respectively. Alignment is at the onset of the first stimulus. Small circles mark neural states at 50 ms intervals. The speed of the neural trajectories is reflected by the distance between consecutive small circles. *(B)* Relative distance, 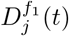, between the *j*-th *f*_1_ trajectory and the reference trajectory for 24Hz, computed during presentation of the first stimulus. Shaded areas represent the mean ± 95% CIs from 100 bootstrap resamples. *(C)* State-space neural trajectories during the first second of the delay period (3 PCs and 44% of variance). Since the first second is common to all delay conditions (1, 1.5, 2, 2.5 and 3 seconds), the analysis of population activity was restricted to this initial interval. Alignment is at the offset of the first stimulus. Small circles represent neural states at 100 ms increments. *(D)* Relative distances computed during the first second of the delay period, 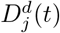.*(E–H)* Same state-space analysis for monkey two. The variance explained in panels *E* and *F* (presentation of *f*_1_) is 76.6%, and in panels *G* and *H* is 58.7%. Panels *G* and *H* are restricted to the first second of the delay, although the delay duration for monkey two was 3 seconds. Only correct trials were considered.

**Fig. 5.**
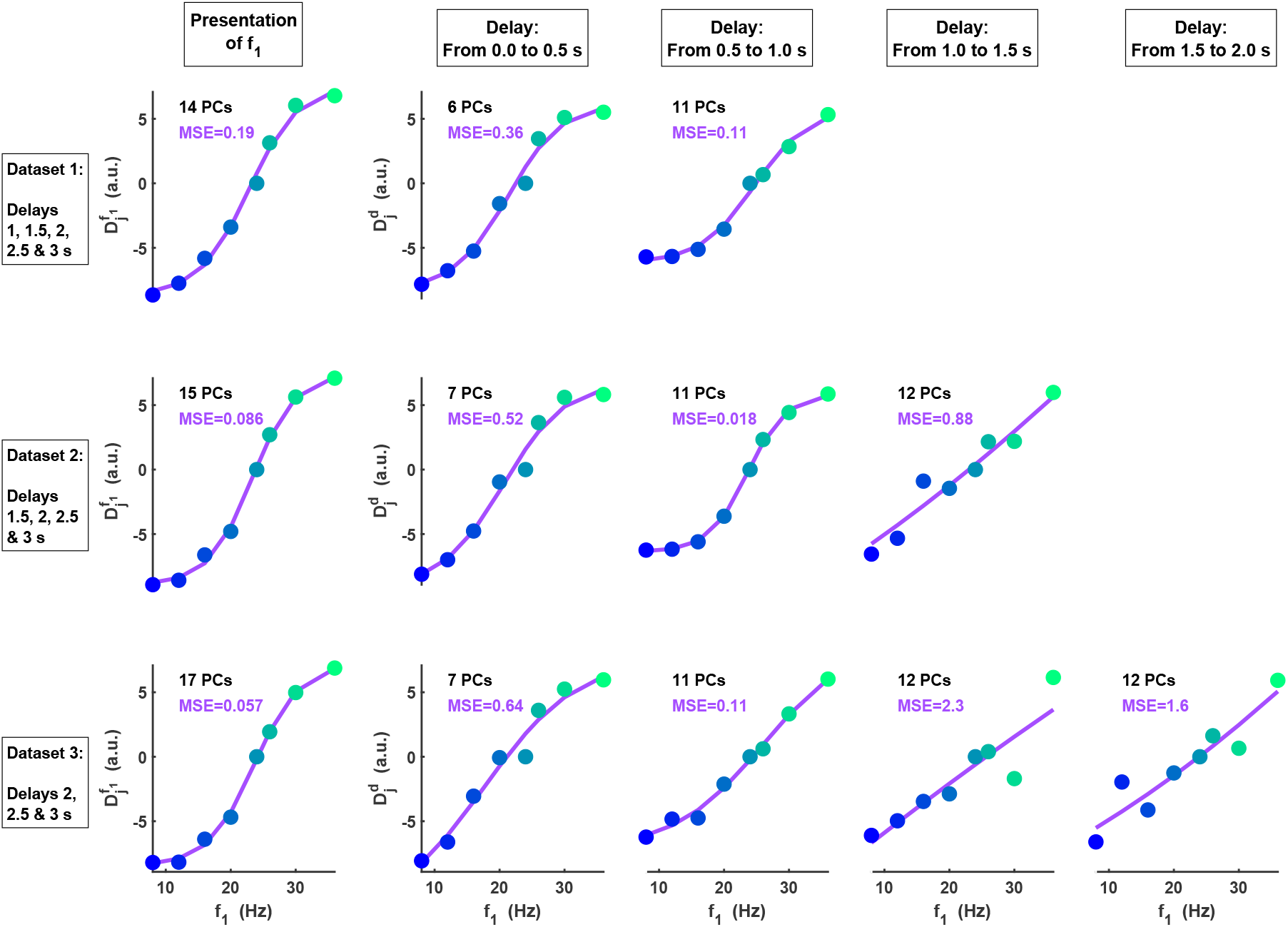
Averaged relative distances for monkey one. Temporal mean of the relative distance, *D*_*j*_, between each trajectory (*j* = 1, …, 8) and the reference trajectory with *f*_1_ = 24Hz. Colored dots represent the averaged distances for individual trajectories, with colors indicating the eight possible *f*_1_ values as in Fig. 4. The purple line is a sigmoidal fit to the data circles; the fit’s mean squared error (MSE) is also indicated in purple. Each column corresponds to a different time segment within the trial: the first column represents the presentation of the first stimulus 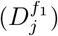, and the subsequent columns represent successive 500 ms intervals of the delay period 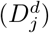.Rows correspond to different datasets used to construct the state-space. The first row includes all delay durations (1, 1.5, 2, 2.5 and 3 seconds). The second row excludes trials with 1s delay, and the third row excludes delays of 1s and 1.5s. As we move down the rows, the dataset becomes progressively smaller, introducing noise unrelated to the delay period. Thus, the first panel of each row serves as a reference to assess the noise from data loss, allowing us to isolate the effects of delay-related noise on the sigmoid from those due to dataset reduction. The state space in each panel was constructed using as many PCs as needed to explain at least 85% of the variance in the presentation of the first stimulus, and at least 60% (61-63%) for each 500 ms segment of the delay period, as indicated in each panel. Only correct trials were considered.

### Relationship Between S2 State-Space Geometry and Behavior

The temporal mean of the relative distances between trajectories as a function of stimulus value exhibits a sigmoidal pattern, both during stimulus presentation and throughout the WM period (Fig. 5). Previous studies have associated this geometric property with behavioral responses, suggesting that the sigmoidal dependence of those neural representations reflects the contraction toward the mean stimulus, or central tendency, observed in animal behavior [7], [23], [24]. More specifically, the Bayesian estimator of the stimulus obtained with a Bayesian behavioral model exhibits a sigmoidal dependence on stimulus frequency and accounts for the relative distances between PFC neural trajectories more accurately than alternative estimators, such as the true stimulus value or the maximum a posteriori (MAP) estimate [7]. This relationship between population neural activity and behavior has also been found in spiking recurrent neural networks trained for delayed response tasks [12].

Drawing from this experience, the analysis in Fig. 5 suggests that the Bayesian estimator, *f*_1,Bayes_, should closely match the relative distances between state-space trajectories. However, for comparison, we also considered two alternative estimators: the true stimulus value (*f*_1_) and the maximum a posteriori (MAP) estimator (*f*_1,MAP_; see Methods, Eqs. 13-14). Note that, as previously done for the Bayesian estimator, the propagated-belief model was adjusted by incorporating the MAP estimator in Eq. 6 (see SI Appendix, Fig. S1 and Table S2). To determine which of them best captured the relative distances between neural trajectories during the stimulus presentation phase, we applied the following procedure. We conducted analyses during three key epochs: the presentation of *f*_1_ and the first two half-second intervals of the delay period. For each epoch, we evaluated the temporal mean of the relative distances (*D*_*j*_) and independently fitted a linear model of the form *aD*_*j*_ + *b* to each of the three estimators. The goodness of fit was assessed by computing the root mean square error (RMSE) (Fig. 6). During the stimulus presentation, the results (Fig. 6*A-C*) strongly supported the hypothesis that the distances between neural trajectories encode a Bayesian estimate of *f*_1_. Similarly, throughout the first two segments of the delay period, our findings (Fig. 6*D-I*) indicated that the neural geometry continued to favor the Bayesian estimator, reinforcing its role in WM representation.

**Fig. 6.**
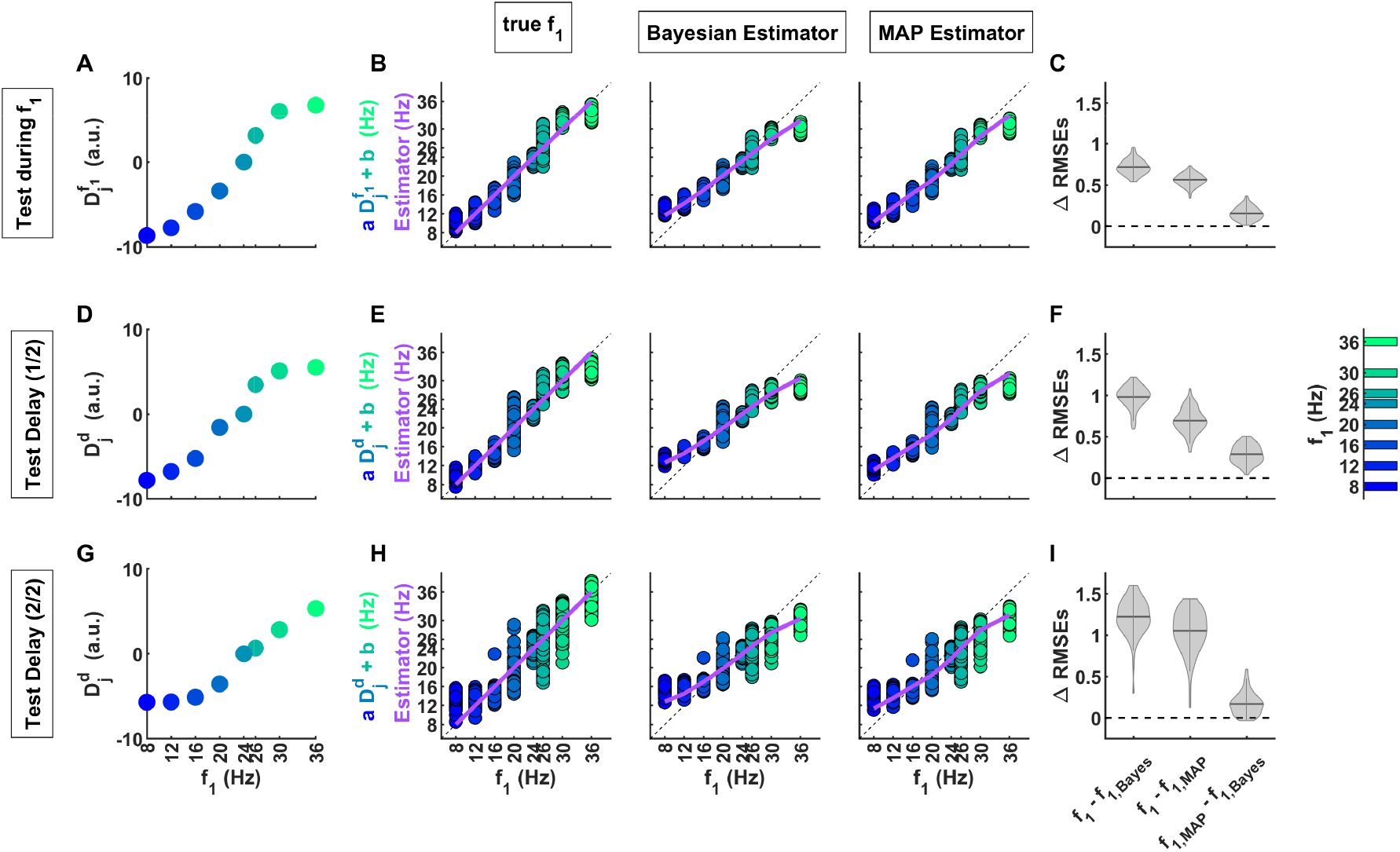
Relative distances between neural trajectories from monkey one code the Bayesian estimator. The following analyses were performed including all delay durations available for monkey one (first row of Fig. 5). *(A-C)* Activity-behavior relationship during the presentation of the first stimulus. *(A)* Temporal mean of relative distances, 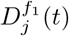, as function of *f*_1_ (Fig. 5, first row, first panel). *(B)* Linear regression of the mean relative distances 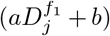 to the true *f*_1_-value *(Left)*, the Bayesian estimator *(Middle)* and the MAP estimator *(Right)* as a function of *f*_1_. Colored dots show regression estimates from 100 bootstrap resamples. Purple solid lines indicate the true *f*_1_ (*Left*), its Bayesian estimator *(Middle)* and its MAP estimator *(Right)*. The unity line is indicated by a black dotted line. *(C)* Violin plots illustrate the empirical probability density of the ΔRMSE values along the y-axis, reflecting the difference in RMSE between pairs of hypotheses (*f*_1_, Bayesian estimator, and MAP estimator) shown on the x-axis. A positive ΔRMSE indicates that the second estimator in the comparison provides a better fit to the geometric feature. Across all tests, the Bayesian estimator consistently outperforms the alternatives. The width of the gray region corresponds to the density of shuffled samples within that range. A horizontal line inside each violin denotes the mean of the distribution. *(D-F)* Activity-behavior relationship during the first 500ms of the delay period (Fig. 5, first row, second panel). Same format as *A-C*. Again, Bayesian hypothesis is chosen by the 100 resamples. *(G-I)* Activity-behavior relationship during the second 500ms segment of the delay period (Fig. 5, first row, third panel). One more time, Bayesian estimator is preferred. The variability across bootstrap resamples increases compared to the test conducted during the first 500ms segment of the delay (panel *E*). This may reflect a progressive loss of neurons with information about *f*_1_ over the course of the delay: since only a subset of the recorded neurons encodes the first stimulus—and this number decreases as the delay progresses—bootstrap resampling with replacement becomes increasingly likely to select non-informative neurons, resulting in greater variability across resamples.

Taken together, the results of these three analyses provide strong evidence that the representation of the first frequency is well described by its Bayesian estimator. This conclusion was further supported by applying the same analysis to the data from monkey two (SI Appendix, Fig. S7). Notably, the population dynamics inherently transformed the actual value of *f*_1_ into its Bayesian estimate, without requiring explicit computations to integrate prior knowledge with sensory input. This implicit transformation was also noticed in our analysis of PFC neural activity in the same task [7]. It is also reminiscent of findings from time interval estimation studies [23], [24] and aligns with observations in simulated recurrent neural networks trained to discriminate between temporal intervals [12].

## Discussion

Understanding how neural activity gives rise to perceptual biases is essential for uncovering the computational principles that shape decision-making. In a previous study, we analyzed prefrontal cortex (PFC) activity during the vibrotactile discrimination task and found that this region not only encoded stimulus information but also exhibited a neural correlate of the contraction bias—where stimuli were represented as shifted toward the center of the stimulus distribution [7]. This distortion was captured by the geometry of the neural state-space trajectories, which conformed closely to a Bayesian estimator, suggesting that PFC population activity embeds a neural correlate of Bayesian inference in perceptual decision-making.

Motivated by these findings and by the growing evidence that working memory (WM) is distributed across multiple cortical areas, in the present study we turned our attention to the secondary somatosensory cortex (S2). S2 has long been implicated in tactile processing and is known to participate in WM-related activity [6]. However, its contribution to perceptual biases has not been investigated. Notably, previous work has shown that stimulus information in S2 neurons fades shortly after the first stimulus offset and re-emerges only upon presentation of the second stimulus [6], [20]. This temporal profile raises the question of whether S2 plays an active role in maintaining a distorted representation of the stimulus during the WM delay period.

We investigated the extent to which the representational mechanisms observed in PFC—specifically, those pointing to a Bayesian origin of the contraction bias—are also present in S2. Our analyses of neural population trajectories suggest that perceptual biases do not emerge exclusively from high-level cortical areas, but rather from a distributed processing architecture that includes early sensory regions. Although WM representations in S2 are less robust and sustained than those in PFC, they nonetheless exhibit key geometric features of the PFC manifold. Notably, we observed a warped encoding of the first stimulus not only during the presentation of the stimulus but also during the mnemonic delay. This suggests that the perceptual bias, and its Bayesian interpretation, may begin to take shape at earlier stages of the cortical processing hierarchy.

These findings raise important questions about the functional contributions of sensory areas to WM and perception: Do these regions merely reflect distortions inherited from downstream areas, or do they actively contribute to shaping a global warped representation of the first stimulus? Although this experiment cannot answer this question, a growing body argues that WM is not localized to a single region but instead relies on interactions across a distributed network of cortical areas [2]–[4]. This interpretation is further supported by recent theoretical models of large-scale cortical dynamics. A recent multiregional model of distributed cognition in the cortex proposed that WM can be sustained not only through local recurrent excitation within individual areas but also through the recurrent loops formed by long-range inter-areal connections [25]–[27]. According to this framework, even when a single area lacks sufficient local excitatory strength to autonomously maintain persistent activity, stable WM representations can still emerge at the network level via interactions across areas. Remarkably, the model predicts a sharp transition in mnemonic capacity across the cortical hierarchy: As one moves along the hierarchy, areas either exhibit robust persistent activity or remain largely transient, with a discontinuity—or bifurcation in space—in firing-rate dynamics marking the boundary between these two regimes. This theoretical insight closely parallels the empirical observations from our task: PFC—a high-level associative area—exhibits robust and sustained WM-related activity, characterized by a warped representation of the stimulus consistent with a Bayesian contribution to the contraction bias. In contrast, S1, situated at the bottom of the cortical hierarchy, shows no evidence of WM representations [5]. Notably, S2, positioned just above S1, displays weak yet organized activity patterns suggestive of an emerging WM representational geometry that resemble that observed in PFC.

Taken together, our results suggest that the contraction bias in perceptual decision-making is embedded within a distributed and hierarchical system in which both higher-order and early sensory areas contribute. Future studies should aim to unravel how these distributed representations interact over time and whether similar mechanisms of distortion and inference extend across other sensory modalities and cortical circuits.

## Methods

### Experimental Data

We analyzed both behavioral data and neural recordings from two monkeys (RR11 and RR13, referred to here as monkey one and monkey two, respectively) [6]. Both monkeys had been trained to perform a somatosensory vibrotactile frequency discrimination task (Fig. 1*A*), each with a distinct set of frequency classes (Fig. 1*B*). While monkey one performed the task with randomly varying delay durations across trials (1, 1.5, 2, 2.5, or 3 seconds), monkey two was recorded performing the task with a fixed 3-second delay period.

Recordings were obtained across multiple experimental sessions for both animals. Each session consisted of approximately 10 blocks, with each block containing one trial per class in the set. Classes were randomly selected without replacement within each block. In the case of monkey one, 15,614 trials were collected across 242 sessions. Behavioral analyses were performed by grouping these trials according to delay duration (trial distribution by delay is shown in SI Appendix, Table S1). Based on a stimulus tuning analysis [6], we selected 335 neurons from monkey one, recorded in 120 sessions and yielded a total of 10,104 trials for the analysis of neural activity. On the other hand, we selected 149 neurons from monkey two, recorded across 42 sessions, yielding a total of 3,736 trials used for both behavioral and neural activity analyses.

### Data analyses: Accuracy curves

To evaluate behavioral performance in both monkeys, we computed the accuracy as the percentage of correct responses as a function of the stimulus class index (Fig. 1*D,E*). For monkey one, separate accuracy curves were constructed for each delay duration using the corresponding subset of trials (SI Appendix, Table S1).

*Phenomenological measure of contraction bias*. To quantify the contraction bias from behavior, we computed a phenomenological measure *B* [7], [12] based on the root-mean-square deviation of the probability of responding *S* ≡ “*f*_1_ *< f*_2_” (i.e., *P* (*S*|*C*)) across all stimulus classes (*C* = 1, …, 10), using classes 3 and 8 as references for the upper and lower diagonals of the stimulus set, respectively (Fig. 1*B*)

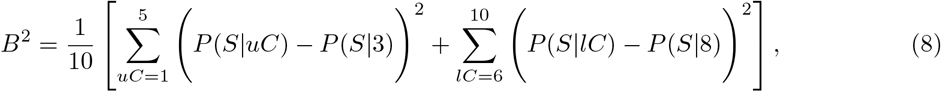

where *uC* and *lC* denote classes on the upper and the lower diagonals of the stimulus set, respectively. A value of *B* = 0 indicates that there is no bias, while the maximum possible bias corresponds to *B*^2^ = 3*/*5.

### Data analyses: Mutual Information

Mutual information (MI) was used to quantify how much information individual neurons conveyed about the first stimulus, *f*_1_, following the original formulation by Shannon [29]. The analysis focused on the response of single neurons, measured as the spike count *r*_*t*_ within the *t*-th time bin. Spike counts were computed using a 250 ms window sliding in 25 ms steps, yielding a set of values *r* = {*r*_1_, …, *r*_*L*_}, where *L* is the total number of bins. The complete collection of these values is denoted by *R*. To evaluate information about the first stimulus, all possible values of *f*_1_ were grouped in the set *F*_1_. Using these definitions, the MU between the neural response and *f*_1_ was computed as [28]

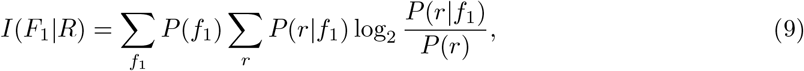

*P* (*f*_1_) is the *f*_*x*_ probability, while *P* (*r* | *f*_1_) denotes the conditional probability of observing a neural response *r* given that *f*_1_ was delivered. The MI in Equation (9) was computed using the MATLAB-based toolbox provided by [30], which also outlines methods to correct for the upward bias that arises when estimating MI from finite data. To address this bias, we applied an analytical correction in combination with a bootstrap procedure. The analytical approach, originally proposed by [31], assumes that the bias can be approximated by a second-order expansion in the inverse of the number of trials, *N*_trial_

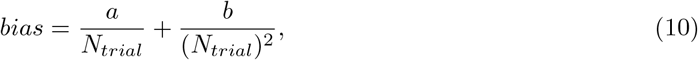

in this formulation, *a* and *b* correspond to the first two terms of the correction series. However, this analytical correction alone was not sufficient, as the series expansion does not fully capture the bias introduced by limited sampling. To further address this limitation, we applied an additional bootstrap-based procedure, also provided in the toolbox. This involved randomly reassigning stimulus labels to neural responses, thereby removing the true association between them. The average MI obtained across these resampled trials was used as an estimate of the residual bias. Instead of subtracting this value from the corrected information, we used it as a baseline to evaluate the statistical significance of the corrected MI (see SI Appendix, Figs. S2, S3 and S4). Significance at each time bin was determined by comparing the corrected MI against this bootstrap-estimated baseline. In particular, we applied a temporal cluster-based criterion: significance required that the MI exceed the baseline threshold for at least three consecutive temporal bins. Shorter sequences of significant bins (one or two bins) were considered spurious and thus excluded from further analysis.

Additionally, we computed the percentage of neurons that exhibited statistically significant MI about *f*_1_ over time (Fig. 3; SI Appendix, Fig. S5). To assess the significance of this proportion, we used a shuffling procedure in which firing rates from all neurons and trials were pooled together and randomly permuted 25 times without replacement. For each time bin, we then calculated the mean and corresponding confidence intervals of the percentage of neurons with significant MI based on the shuffled data. These values are shown as horizontal shaded bands in Fig. 3 and SI Appendix, Fig. S5.

### Data analyses: Firing Rates

Trial-averaged firing rates were estimated by conditioning on *f*_1_. For both monkeys, a temporal window of 100 ms was used, sliding in 10 ms steps, during the analysis of the first stimulus presentation (aligned to its onset) and the delay period (aligned to the offset of *f*_1_). An exception was made for the analyses during the delay period shown in SI Appendix, Figs. S6 and S7, which focus on the delay activity of monkey two across three consecutive 1000 ms segments. In this case, firing rates were computed using a 200 ms window sliding every 20 ms.

### Data analyses: State-space Analysis and Geometry

*Principal Component Analysis*. To identify low-dimensional neural state spaces, we applied principal component (PC) analysis to the population firing rates averaged across trials and conditioned on *f*_1_. For visualization purposes (Fig. 4), trajectories were projected onto the first three PCs, capturing 66% and 50% of the variance during the *f*_1_ presentation and the first second of the delay, respectively, in monkey one, and 76% and 59% in monkey two. In contrast, for the quantitative analysis of trajectory geometry (Fig. 5; SI Appendix, Fig. S6), we constructed PC spaces that explained at least 85% of the variance during the *f*_1_ presentation, and at least 60% during the delay period. PCA was performed independently within each delay segment—500 ms windows for monkey one and 1000 ms windows for monkey two.

*KiNeT Method. Relative distances between state-space trajectories*. To quantify the separation between neural trajectories, we used the Kinematic analysis of Neural Trajectories (KiNeT) approach [24]. This method compares a given trajectory *j* to a selected reference trajectory *i* by measuring their relative distances over time. At each time point *t*, we identified the neural state of the reference trajectory *s*_ref_(*t*), and searched for the closest state *s*_*j*_(*t*) on trajectory *j* in terms of Euclidean distance. The distance between these two points defined the relative distance *D*_*j*_(*t*) at that time (*j* = 1, …, *N*_1_, with *N*_1_ = 8 for monkey one and *N*_1_ = 7 for monkey two). For monkey one, the reference trajectory corresponded to *f*_1_ = 24 Hz, while for monkey two it was set to *f*_1_ = 22 Hz. Relative distances were computed within the PC spaces described above (see *Principal Component Analysis* section). To estimate variability, we performed bootstrap resampling with replacement (100 resamples). For each resample, PC and distance analyses were repeated, and the final estimates were reported as the mean ± 95% confidence intervals.

Finally, to characterize the geometry of the trajectories, we computed the temporal mean, *D*_*j*_, of the relative distances across the presentation of the first stimulus 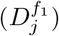 and across successive intervals (500/1000 ms for monkeys one/two, respectively) of the delay period (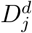; see Fig. 5 and SI Appendix, Fig. S6).

### Bayesian and MAP estimators

In the normative model, propagating the belief throughout the delay period (Eq. 6) requires formulating a hypothesis about how the first stimulus is represented during that interval. In particular, we considered a Bayesian estimator (*f*_1,Bayes_) and a maximum a posteriori (MAP) estimator (*f*_1,MAP_) of the first stimulus. To estimate these inferred representations, we sampled, for each value of *f*_1_, 5,000 noisy observations drawn from the likelihood function *P* (*O*_1_ | *f*_1,estim_, *t*), where *f*_1,estim_ denotes either *f*_1,Bayes_ or *f*_1,MAP_. These simulated observations were then discretized and matched to a predefined observation space ranging from 1 to 100 Hz in 1 Hz steps. For each simulated observation, *o*_*j*_, the Bayesian estimate was computed as the posterior-weighted mean of the possible *f*_1_ values

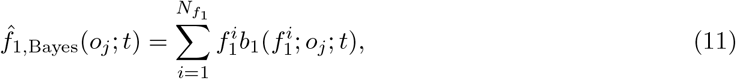

where 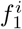 denotes the *i*-th value of the stimulus *f*_1_, *N*_1_ is the number of possible *f*_1_ values (8 for monkey one and 7 for monkey two), and 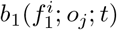 is the belief associated with observation *o*_*j*_ at time *t* (Eq. 6). The overall Bayesian estimate for a given true stimulus value is the average across all simulated observations

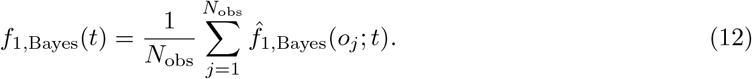

The MAP estimator is the value of *f*_1_ that maximizes the posterior at time *t*

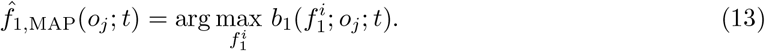

The overall MAP estimate is obtained by averaging the selected values across all observations

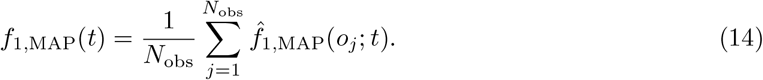

These estimators were used to update the belief about *f*_1_ throughout the delay period. For this reason, estimators were computed at the end of *f*_1_ (that is, the start of propagation) and at subsequent time points: every 500 ms for each delay duration in monkey one, and every 1000 ms in monkey two.

### Relationship between *f*_1_ estimation and state-space geometry

To assess whether the geometry of population neural activity encoded an estimate of the first stimulus, we confronted three estimators: the true *f*_1_, the Bayesian estimator *f*_1,Bayes_, and the MAP estimator *f*_1,MAP_.

It should be noted that distinct Bayesian and MAP estimators were obtained for each of the five delay durations in monkey one, as the accuracy curves were fitted separately (see Fig. 2 and SI Appendix, Fig. S1). For this reason, we combined the estimators across all delay durations to compute an averaged (Bayesian/MAP) estimator, weighting each by the fraction of trials corresponding to that delay in the dataset used to compute the relative distances (see SI Appendix, Table S1).

To determine which estimator best explained the temporal mean of the relative distances between trajectories (*D*_*j*_), we compared the three hypotheses—true *f*_1_, Bayesian, and MAP estimators—during both the presentation of *f*_1_ and the delay period (Fig. 6)

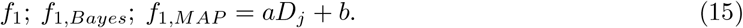

To evaluate the quality of the fits, we compared their Root Mean Square Errors (see Figs. 6*C, F,I* for monkey one and SI Appendix, Fig. S7*C,F,I* for monkey two). Statistical significance was assessed via a bootstrap procedure: 100 resamples were generated by randomly sampling neurons with replacement from the original population. Each test was then repeated on the resampled neural populations, and the resulting variability was quantified.

## Code and Data Availability

Matlab codes, as well as the raw data necessary for full replication of the figures will be available upon publication.

## Acknowledgements

We gratefully acknowledge Ignacio Fresneda Gallardo, Jorge Manrique Ruiz and Fernando Plaza Munõz for their valuable contributions to the preprocessing of the experimental data.

## Author contributions

N.P. and R.R. designed research. L.S.-F. implemented the numerical computations and analyzed the data. L.S.-F. and N.P. wrote the paper. R.R. revised the text.

