## Supplementary material for "Early Emergence of Perceptual Biases in Working Memory": SI Appendix

### Supporting Information for
### Early Emergence of Perceptual Biases in Working Memory.

Luis Serrano-Fernández[a,b], Ranulfo Romo[c] and Néstor Parga[a,b]

[a] Departamento de Física Teórica, Universidad Autónoma de Madrid, 28049 Madrid, Spain.
[b] Centro de Investigación Avanzada en Física Fundamental, Universidad Autónoma de Madrid, 28049 Madrid, Spain.
[c] El Colegio Nacional, 06020 México DF, México.

This PDF file includes:

Figs. S1 to S7

Tables S1 to S2

### Supplementary Figures

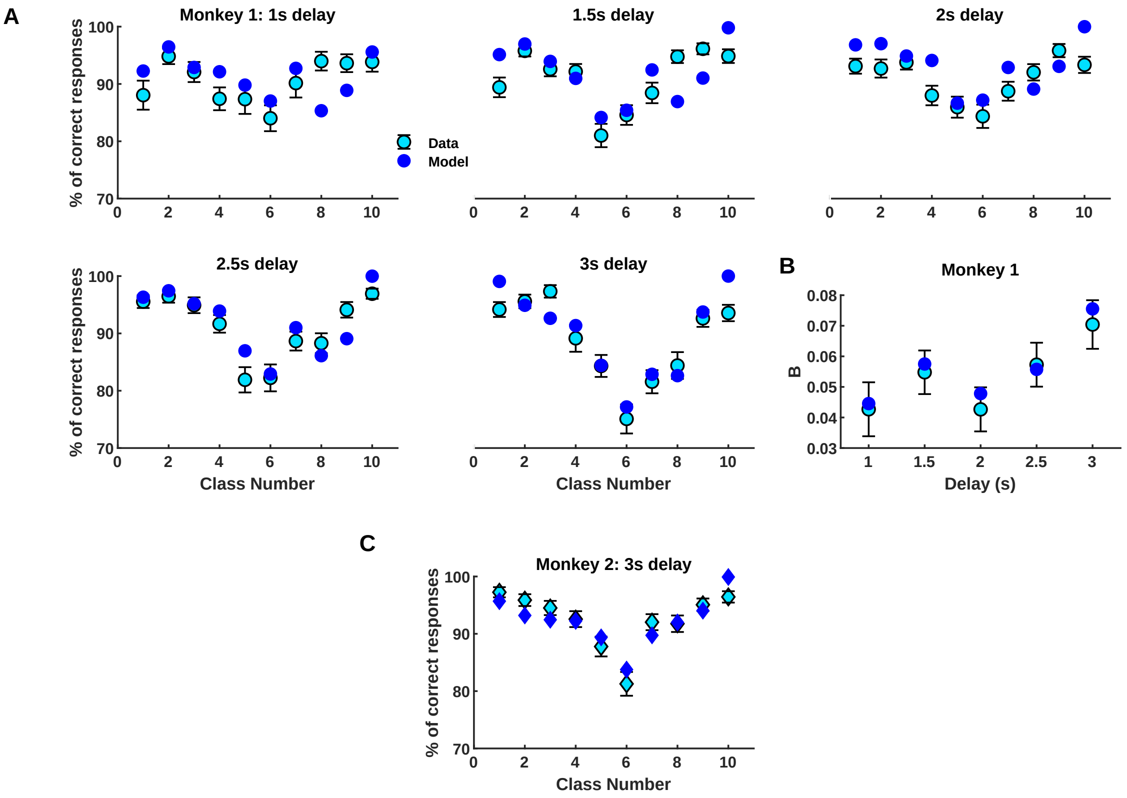


**Fig. S1. MAP fits of the accuracy curves**. *(A)* Fits for monkey one. Each panel corresponds to one of the five possible delay durations (1s, 1.5s, 2s, 2.5s and 3s). Cyan circles with error bars correspond to behavioral data, and dark blue circles represent the MAP fit. *(B)* Contraction-bias measure ($B$; see Eq. 8 in Materials and Methods) obtained from behavioral performance (cyan markers) and the MAP fit (dark blue markers). *(C)* MAP fit for monkey two. Cyan diamonds with error bars correspond to behavioral data, and dark blue diamonds represent the Bayesian fit. Error bars computed with 1,000 bootstrap resamples.

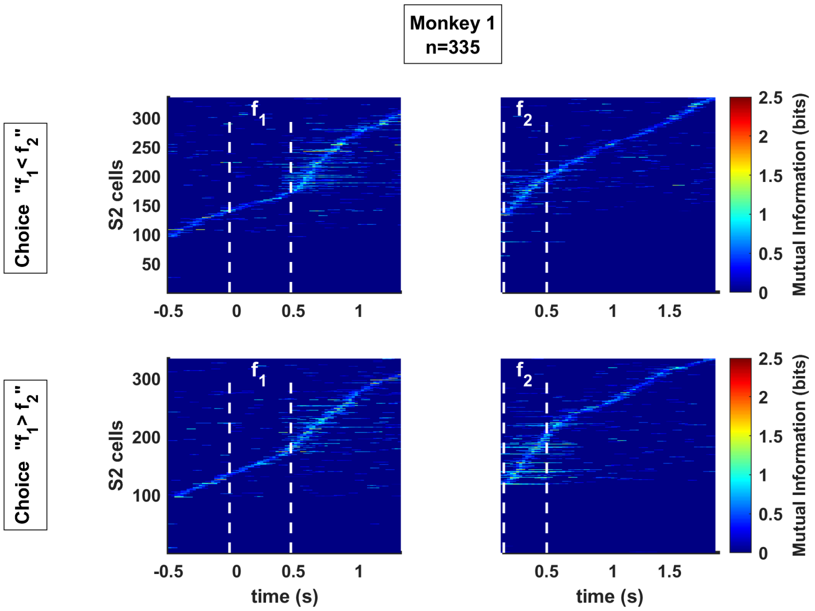


**Fig. S2. Mutual information (MI) about the current-trial $f_1$ in monkey one**. MI over time computed from 335 neurons recorded in monkey one, conditioned on rewarded current trials with a fixed choice: "$f_1 < f_2$" (*Top*) or "$f_1 > f_2$" (*Bottom*). This constraint effectively fixes the stimulus class in the current trial. Non-significant MI values are shown in dark blue (see Methods). Neural recordings from all delay conditions were pooled to analyze MI, aligning activity to the onset of the first stimulus (left panels) or the second stimulus (right panels).

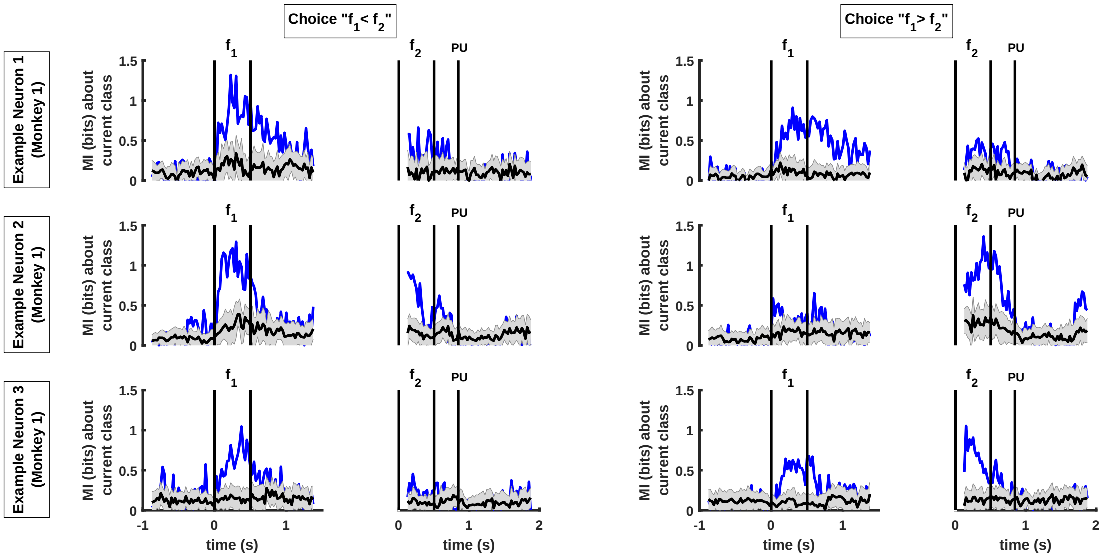


**Fig. S3. Mutual information (MI) about the current-trial $f_1$ in three example single neurons from monkey one**. Time-resolved MI computed for three neurons (one per row) recorded in monkey one, conditioned on rewarded current trials with a fixed choice: "$f_1 < f_2$" (*Left*) or "$f_1 > f_2$" (*Right*). This constraint effectively fixes the stimulus class in the current trial. Horizontal black shadings indicate the mean and standard deviation obtained from shuffled data (see Methods). Neural recordings from all delay conditions were pooled to analyze MI.

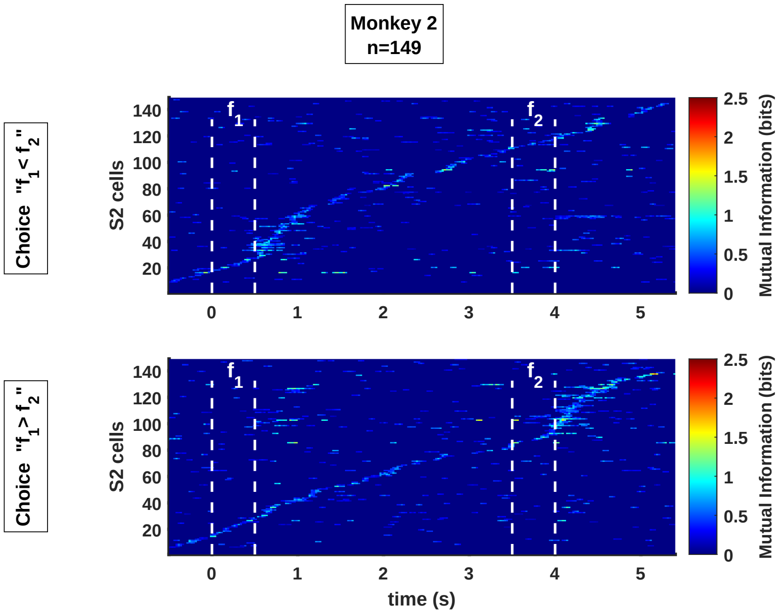


**Fig. S4. Mutual information (MI) about the current-trial $f_1$ in monkey two**. MI over time computed from 149 neurons recorded in monkey two, conditioned on rewarded current trials with a fixed choice: "$f_1 < f_2$" (*Top*) or "$f_1 > f_2$" (*Bottom*). This constraint effectively fixes the stimulus class in the current trial. Non-significant MI values are shown in dark blue (see Methods).

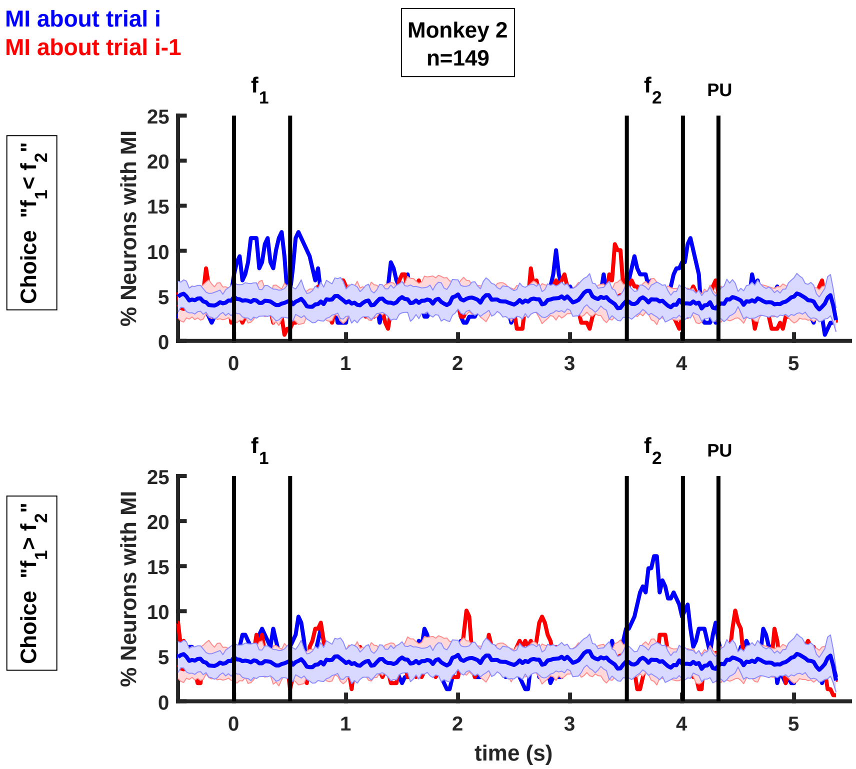


**Fig. S5. S2 neurons from monkey two exhibit significant mutual information (MI) about current-trial $f_1$, but not about the previous-trial $f_1$.** A small percentage of neurons showing significant MI with respect to the first stimulus of the current trial (blue line) versus that of the previous trial (red line). The analysis was restricted to correct trials in which the current choice was fixed (*Top*: "$f_1 < f_2$"; *Bottom*: "$f_1 > f_2$"). Note that this constraint effectively fixes the stimulus class in the current trial. Horizontal blue and red shadings indicate the mean and standard deviation computed from shuffled data for the current and previous trial, respectively (see Methods).

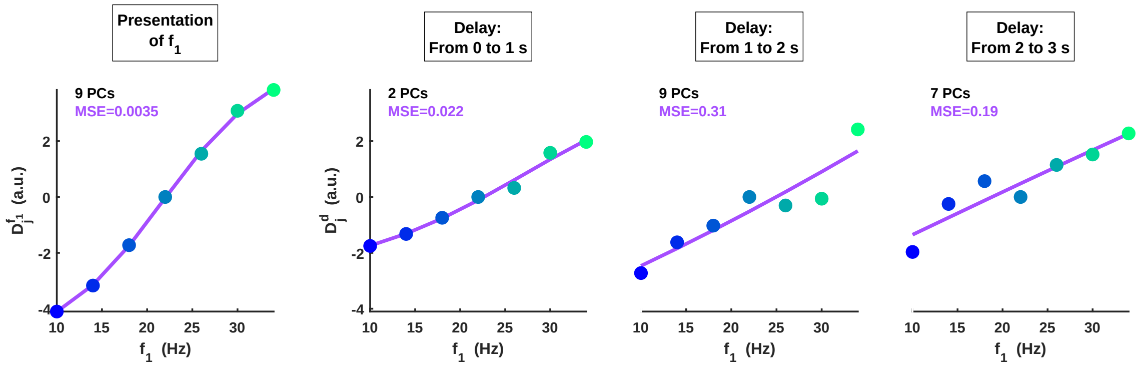


**Fig. S6. Averaged relative distances for monkey two**. Temporal mean of the relative distance, $D_j$, between each trajectory ($j = 1, \ldots, 7$) and the reference trajectory with $f_1 = 22$Hz. Colored dots represent the averaged distances for individual trajectories, with colors indicating the eight possible $f_1$ values. The purple line shows a sigmoidal fit to the data circles; the fit's mean squared error (MSE) is also indicated in purple. Each panel corresponds to a different time segment within the trial: the first panel represents the presentation of the first stimulus ($D_j^{f_1}$), and the subsequent panels represent 1000ms intervals of the delay period ($D_j^d$). The state-space in each panel was constructed using as many PCs as needed to explain at least 85% of the variance in the presentation of the first stimulus, and at least 60% for each 1000ms segment of the delay period. Only correct trials were considered.

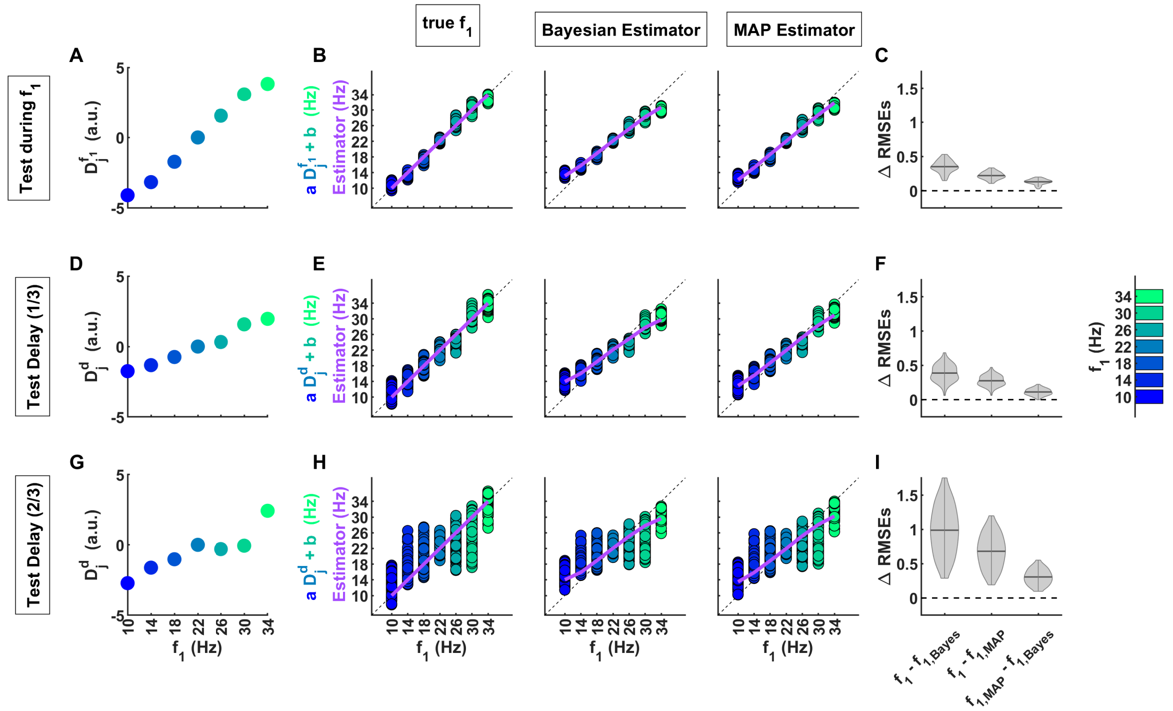


**Fig. S7. Relative distances between neural trajectories from monkey two code the Bayesian estimator**. *(A-C)* Activity-behavior relationship during the presentation of the first stimulus. *(A)* Temporal mean of relative distances, $D_j^{f_1}(t)$, as function of $f_1$ (Supplementary Fig. S6, first panel). *(B)* Linear regression of the mean relative distances ($aD_j^{f_1} + b$) to the true $f_1$-value *(Left)*, the Bayesian estimator *(Middle)* and the MAP estimator *(Right)* as a function of $f_1$. Colored dots show regression estimates from 100 bootstrap resamples. Purple solid lines indicate the true $f_1$ *(Left)*, its Bayesian estimator *(Middle)* and its MAP estimator *(Right)*. The unity line is indicated by a black dotted line. *(C)* Violin plots illustrate the empirical probability density of the $\Delta$RMSE values along the y-axis, reflecting the difference in RMSE between pairs of hypotheses ($f_1$, Bayesian estimator, and MAP estimator) shown on the x-axis. A positive $\Delta$RMSE indicates that the second estimator in the comparison provides a better fit to the geometric feature. The width of the gray region corresponds to the density of shuffled samples within that range. A horizontal line inside each violin denotes the mean of the distribution. *(D-F)* Activity-behavior relationship during the first 1000ms of the delay period (Supplementary Fig. S6, second panel). Same format as *A-C*. *(G-I)* Activity-behavior relationship during the second 1000ms segment of the delay period (Supplementary Fig. S6, third panel).

### Supplementary Tables

**Table S1. Number of trials per delay condition for monkey one.** The table shows the number and percentage of trials for each delay duration (1, 1.5, 2, 2.5, and 3 seconds), separately for behavioral and neural activity analyses. Each analysis used a different dataset: for behavioral analyses, all trials from 242 recording sessions were included; for neural activity analyses, neurons were filtered based on their tuning properties, which reduced the dataset to 120 sessions and, consequently, decreased the number of available trials per delay condition.

| | **Behavior analyses # trials (%)** | **Neural act. analyses # trials (%)** |
|---|---|---|
| 1s delay | 2168 (13.89%) | 1591 (15.75%) |
| 1.5s delay | 3846 (24.63%) | 2542 (25.16%) |
| 2s delay | 3429 (21.96%) | 2244 (22.21%) |
| 2.5s delay | 3179 (20.36%) | 1856 (18.37%) |
| 3s delay | 2992 (19.16%) | 1871 (18.51%) |
| Total Trials | 15614 | 10104 |

**Table S2. MAP fits across delay conditions for two monkeys**. The fitted behavioral datasets correspond to the five delay conditions tested in monkey one (1, 1.5, 2, 2.5 and 3 seconds), and the 3-second delay condition in monkey two. The second and third columns display the fitted measurement uncertainty parameters ($W_1$ and $W_2$). The fourth column shows the fitted $\eta$ parameter, and the last column reports the root mean square error (RMSE) between behavioral and model performance.

| **Delay (Monkey)** | $W_1$ | $W_2$ | $\eta$ | **RMSE** |
|---|---|---|---|---|
| 1s (M1) | 0.101 | 0.156 | 0.420 | 0.0386 |
| 1.5s (M1) | 0.109 | 0.097 | 0.465 | 0.0399 |
| 2s (M1) | 0.118 | 0.076 | 0.447 | 0.0370 |
| 2.5s (M1) | 0.142 | 0.075 | 0.455 | 0.0172 |
| 3s (M1) | 0.274 | 0.055 | 0.282 | 0.0320 |
| 3s (M2) | 0.050 | 0.075 | 0.514 | 0.0172 |
